# Regulation of retinal neurogenesis by somatostatin signaling

**DOI:** 10.1101/2020.09.26.314104

**Authors:** Kurt Weir, Dong Won Kim, Seth Blackshaw

## Abstract

Neuropeptides have been reported to regulate progenitor proliferation and neurogenesis in the central nervous system. However, these studies have typically been conducted using pharmacological agents in *ex vivo* preparations, and *in vivo* evidence for their developmental function is generally lacking. Recent scRNA-Seq studies have identified multiple neuropeptides and their receptors as being selectively expressed in neurogenic progenitors of the embryonic mouse and human retina. This includes Sstr2, whose ligand somatostatin is transiently expressed by immature retinal ganglion cells. By analyzing retinal explants treated with selective ligands that target these receptors, we found that Sstr2-dependent somatostatin signaling induces a dose-dependent inhibition of photoreceptor generation while increasing the relative fraction of primary progenitor cells. These effects were confirmed by scRNA-Seq analysis of retinal explants and abolished in *Sstr2*-deficient retinas. Although no changes in the relative fraction of primary progenitors or photoreceptor precursors were observed in *Sstr2*-deficient retinas *in vivo*, scRNA-Seq analysis demonstrated accelerated differentiation of neurogenic progenitors. We conclude that Sstr2 signaling may act to negatively regulate retinal neurogenesis in combination with other retinal ganglion cell-derived secreted factors such as Shh, although *in vivo* Sstr2 is dispensable for normal retinal development.

## Introduction

Neurotransmitters play important roles in regulating neurogenesis and development. Although neurotransmitters are generally considered in terms of neuronal signaling, neurotransmitter expression precedes the nervous system on both evolutionary and developmental time scales^1^. In fact, neurotransmitter genes are found in species without a central nervous system like sponges and the social amoeba *Dictysotelium*^2^. The receptors for several neurotransmitters such as GABA, glutamate, and serotonin are expressed on progenitor cells of the central nervous system prior to the formation of functional synapses^3–5^. The neurotransmitters themselves likely act as extrinsic signaling factors to guide neural development^1,3–5^.

It may be that developmental signaling represents the ancestral role for neurotransmitters before their use in signaling by mature neurons^2^. Neurotransmitters and neurotransmitter receptors are expressed in neural progenitors and/or immature neurons in many different regions of the vertebrate CNS^6–8^. Over the past three decades, there has also been a large number of studies in which pharmacological manipulation of a number of neurotransmitter systems have been reported to modulate multiple different aspects of neuronal development in different brain regions, including progenitor proliferation, neurogenesis, axonal guidance, synaptogenesis and neuronal survival^3,9,10^. The hippocampus is the brain region with the greatest number of reported neurotransmitter developmental signals: ATP, serotonin, VGF, vasoactive intestinal peptide (VIP), histamine, glycine, acetylcholine, dopamine, cholecystokinin, oxytocin, corticotropin-releasing hormone (CRH), ghrelin, nitric oxide (NO), GABA, and glutamate have all been found to play different roles in developmental and/or adult rodent hippocampal neurogenesis^3,6,11–22^. Other regions have fewer reported neurotransmitter developmental signals, like GABA, glutamate, histamine, VIP, and taurine^3,9,13,15^ in neocortical development, and pituitary adenylate cyclase-activating peptide (PACAP) in cerebellar development. Serotonin regulates the development of multiple brain regions. However, the specificity of these ligands -- particularly at the high concentrations often used in these studies -- is unclear. Although mutants for many genes directly related to neurotransmission are available, developmental phenotypes are seldom characterized. The precise role of neurotransmitters in regulating brain development thus remains largely unclear.

The retina is an accessible and relatively less complex CNS region that serves as a useful model for understanding molecular mechanisms controlling the development of more complex brain regions^23^. Extrinsic signaling also plays a critical role in regulating retinal development. In addition to classical growth and differentiation factors that are present in the developing retina -- such as bFGF and TGF alpha^24^, GDF11^25^, VEGF^26^ and SHH^27^ -- a number of neurotransmitters are expressed by retinal progenitors and/or immature retinal neurons^28^. Pharmacological manipulation of serotonin, glutamate, or GABA signaling impacts retinal development in *Xenopus laevis*^1^, and endogenous opiate-like peptides have been reported to regulate neurogenesis in rodent retina^27,29^. However, genetic data to support these findings is lacking. Moreover, until recently, comprehensive and quantitative analysis of the cellular expression patterns of neurotransmitter biosynthetic enzymes and receptors during retinal development has not been available. This has changed with the generation of several large-scale single-cell RNA-sequencing (scRNA-Seq) datasets from developing mouse and human retina^30,31^.

This work has identified four different neuropeptides and/or their receptors as being strongly and selectively expressed during early stages of retina development -- somatostatin, neuropeptide Y, galanin, and proenkephalin. Each of these neuropeptides has been reported to regulate brain development. Somatostatin acts as a generalized growth factor inhibitor, and somatostatin expression in the hypothalamus decreases with age along with hypothalamic neurogenesis^13,32^. Neuropeptide Y promotes proliferation in the subventricular zone and hippocampal precursor cells^6,13,15,33^. It also promotes proliferation of dissociated rat retinal progenitor cells, apparently through NO^34^. Galanin, acting through its receptor GalR3, promotes neurogenesis in the adult hippocampus^6,13,17,35^. Finally, proenkephalin impacts cerebellar development and adult hippocampal neurogenesis^36,37^.

To define any of these neuropeptides as an endogenous regulator of retinal development, we need to identify 1) a source of the ligand, 2) impacts on neurogenesis from activation, and knockout of the appropriate receptors, and 3) an *in vivo* phenotype ^38^. To accomplish this, we use publicly-available single-cell RNA-seq data of mouse and human retinal development^30,31^, a pharmacologic screen on murine embryonic retinal explants, and single-cell sequencing of retinal explants and mutant retinas. We find that though they play roles in development elsewhere in the CNS or in other *in vitro* situations, neuropeptide Y, galanin, and proenkephalin do not impact neurogenesis in embryonic retinal explants. Activation of the somatostatin receptor *Sstr2* reduces the production of photoreceptors in this model. However, somatostatin likely plays a redundant role in retinal development as its knockout does not produce the same effect *in vivo*.

## Results

### Interrogation of scRNA-seq datasets identifies candidate developmental signals

A previously published single-cell RNA sequencing (scRNA-seq) study profiled mouse retinal development across ten-time points. We reasoned that this gene expression dataset could be used to identify candidate neurotransmitters impacting mouse retinal development. We downloaded the dataset and used the stored cell type and age data to investigate the expression of neuropeptides in mouse development.

Cellular expression patterns of neuropeptides are readily identified using scRNA-seq. The neuropeptides somatostatin (Sst), galanin (Gal), neuropeptide Y (Npy), and proenkephalin (Penk) were selected for further study as they showed the greatest correlation of expression with a single cell type (Fig 1A). *Sst* and *Gal* are expressed by mouse retinal ganglion cells most strongly between E14 and P0 (Fig 1A, C, S1A), and by human retinal ganglion cells between gestational day 42 and gestational week 13 (Fig S1D, F). Gal is also strongly expressed in early-stage neurogenic progenitors^39^. *Npy* is expressed primarily by late-stage primary retinal progenitor cells between E18 and P2, and *Penk* is expressed by neurogenic progenitor cells mostly between E14 and P2 (Fig 1A, S1B, C). The expression of neuropeptides by specific cell types over delimited windows during retinal development suggests that they may be acting as signaling molecules impacting retinogenesis.

**Figure 1.**
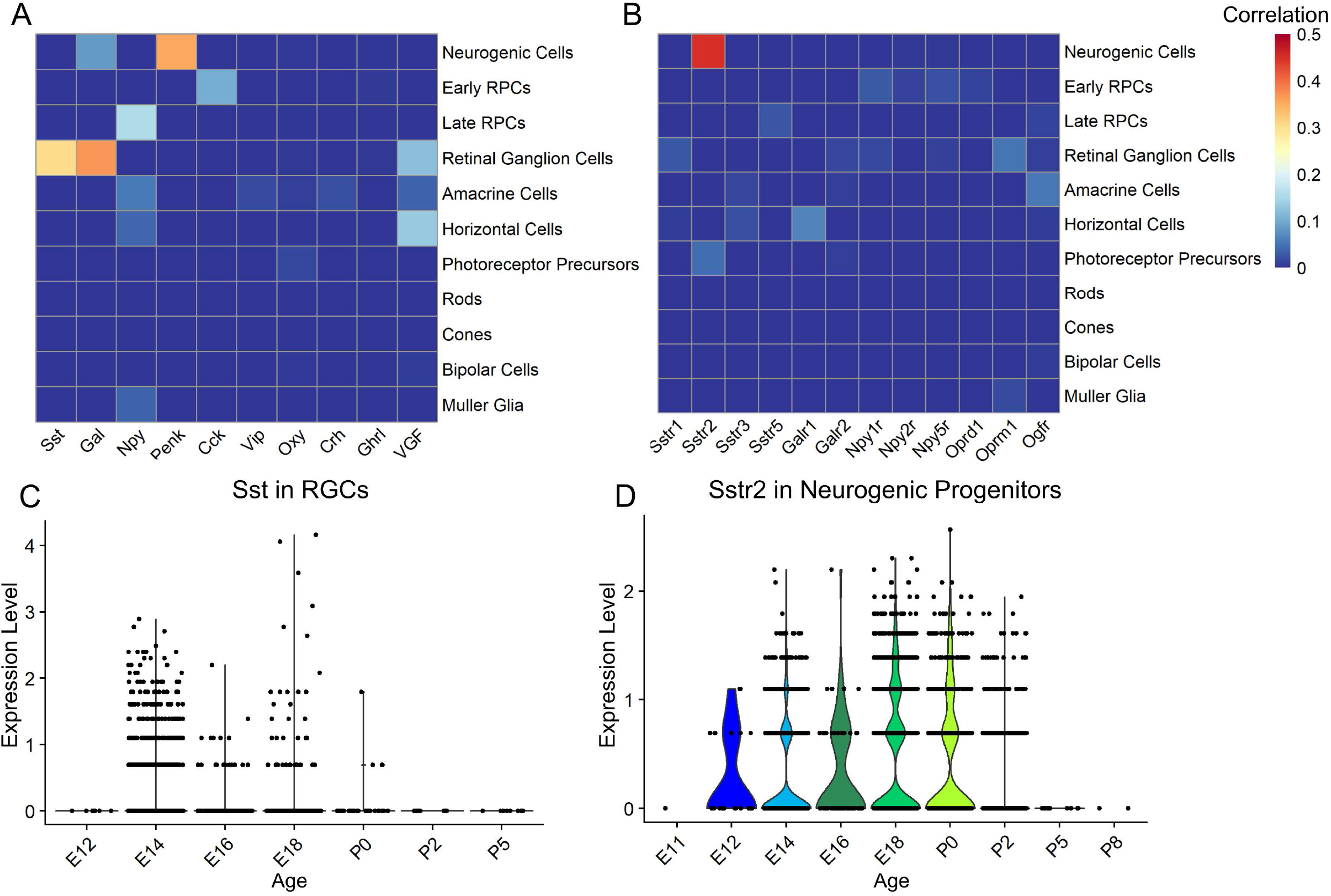
Cell-specific expression of neuropeptides and neuropeptide receptors in developing mouse retina. (A) Heatmap of the correlation of expression of a neuropeptide with a given cell type in mouse retinal development. (B) Correlation of expression of a neuropeptide receptor with a given cell type in mouse retinal development. Sstr4, Galr3, and Npy4r were not expressed at detectable levels in the dataset. (C) Violin plot of Sst expression in retinal ganglion cells across mouse development. Each point represents a single cell. Expression determined by the SCT method in Seurat. (D) Violin plot of Sstr2 expression in neurogenic progenitor cells across mouse development. Correlation calculated using base R function ‘cor()’.

In sharp contrast, with one exception, the receptors for Sst, Gal, Npy, and Penk were either not detected or only barely detectable in both developing mouse and human retina. The one notable exception was somatostatin receptor 2 (*Sstr2*) which is strongly and selectively expressed by neurogenic RPCs from E12 to P2 in mice (Fig 1B, D). This matches the period when *Sst* is prominently expressed in immature RGCs. *SSTR2* is also expressed by neurogenic RPCs in the human retina, while *SST* is also expressed in immature RGCs (Figure S1E, F, G). This led us to hypothesize that Sst could be acting as a signal released by retinal ganglion cells to influence neurogenic progenitors through Sstr2.

### Sstr2 signaling limits the expression of photoreceptor marker genes *in vitro*

We decided to use a pharmacologic screen, treating mouse embryonic retinal explants with small molecule agonists and antagonists to the receptors of the four neuropeptides of interest, as a cost-effective test for an effect on retinal development. Because *Sstr2* was highly and selectively expressed in neurogenic RPCs, while other Sst receptors were not expressed or barely detectable, we used an Sstr2-specific agonist and antagonist. For the other candidates, we ordered the neuropeptides themselves and antagonists general to all of their receptors; except Npy, for which no general antagonist was available. The explants were cultured for a period roughly equivalent to the expression of the relevant neuropeptide *in vivo* with either agonist, antagonist, or solvent control. We used qRT-PCR to measure the relative levels of multiple cell type marker genes between treated retinas and control as a readout of the effects of the treatments on the expression of cell type-specific markers.

Treatment with Npy, galanin, and the galanin receptor antagonist M40 did not produce significant changes in expression for the tested marker genes (Fig 2A, B). Treatment with Met5-enkephalin, a bioactive processed product of Penk, decreased expression of the neurogenic progenitor markers *Atoh7, Neurog2*, and *Sstr2* while the broad-spectrum opiate receptor antagonist naltrexone increased expression for *Atoh7* and *Sstr2* as measured by ANOVA (p-value < 0.0045, Fig 2C). These results suggest that Penk signaling may reduce the proportion of neurogenic progenitor cells in retinal explants. Meanwhile, treatment with the Sstr2-specific agonist (1R,1’S,3’R/1R,1’R,3’S)- L-054,264 consistently nominally significantly decreased expression of the photoreceptor markers *Crx* and *Otx2* (p-value < 0.05, Fig 2D, S2B). This similarly suggests that Sst signaling through Sstr2 negatively regulates photoreceptor differentiation.

**Figure 2.**
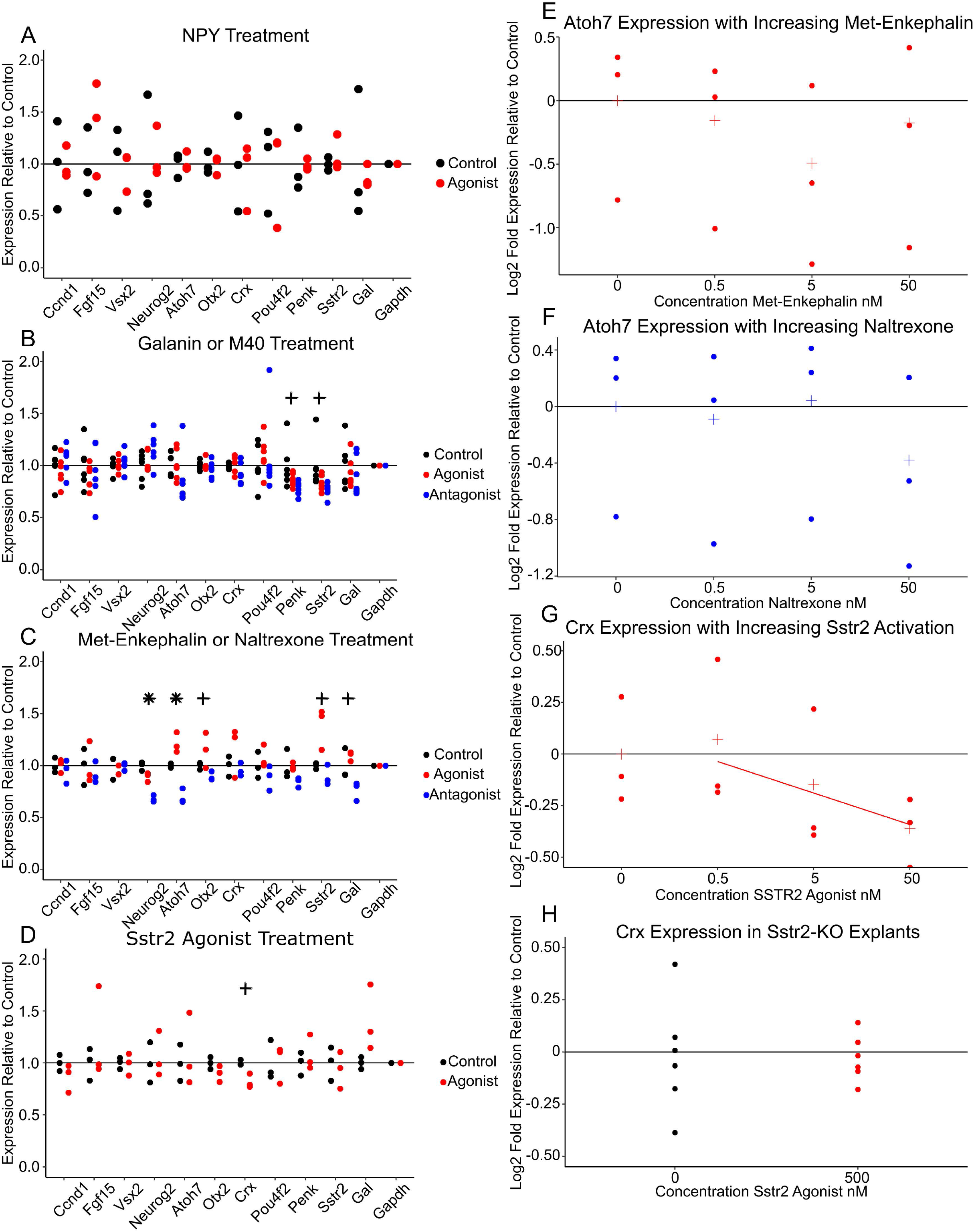
Analysis of cell type-specific markers by qRT-PCR in embryonic retinal explants treated with neuropeptide agonists and antagonists. Cell type marker genes tested include *Ccnd1* (all progenitors); *Fgf15* and *Vsx2* (primary progenitors); *Neurog2* and *Atoh7* (neurogenic progenitors); *Otx2* and *Crx* (photoreceptor precursors); *Pou4f2* (retinal ganglion cells); *Penk, Sstr2*, and *Gal* (neuropeptides in the study, with each of these also being selective markers of neurogenic progenitors); and *Gapdh* (internal control). Each point represents a sample (average of 3 technical replicates). Each sample was tested for each gene. (A) Marker gene expression relative to average control with Npy treatment. Concentration=1 uM. N=3 each condition. Explants treated E18-P2. (B) Marker gene expression relative to average control with Galanin or M40 treatment. Concentration=1 uM. N=6 each condition. Explants treated E14-P0. (C) Marker gene expression relative to average control with Met-Enkephalin or Naltrexone treatment. Concentration=500nM. N= 3 each condition. Explants treated E14-P0. (D) Marker gene expression relative to average control with treatment with SSTR2 agonist (1R,1’S,3’R/1R,1’R,3’S)-L-054,264. Concentration: 500 nM. N=3 each condition. Explants treated E14-P0. (E) Atoh7 expression relative to average control of explants treated with increasing concentration of Met-Enkephalin. N=3 each condition. Explants treated E14-P0. (F) *Atoh7* expression relative to average control of explants treated with increasing concentrations of naltrexone. N=3 each condition. Explants treated E14-P0. (G) *Crx* Expression relative to average control of explants treated with increasing concentrations of Sstr2 agonist. Fitted line added for emphasis. N=3 each condition. Explants treated from E14-P0. (H) *Crx* expression relative to controls in *Sstr2* -/- explants treated with 500 nM Sstr2 agonists. N=6 each condition. Explants treated E14-P0. P-value calculated using ANOVA. + indicates a nominally significant p-value below 0.05. * indicates a p-value significant at a Bonferroni correction below 0.0045.

We next conducted further studies of the potential role of Penk and Sst/Sstr2 in regulating retinal development. To increase our ability to detect an impact on retinal development, the initial assay had been performed using concentrations orders of magnitude in excess of the ligand’s reported EC50 or IC50 values. It is therefore possible that the observed effects were due to nonspecific or off-target effects of these ligands, rather than selective modulation of neuropeptide receptors. In order to test this, we cultured explants in a range of treatment concentrations below that tested previously. We then tested for expression of *Atoh7, Neurog2*, and *Sstr2* following Met5-enkephalin and naltrexone treatment, and for *Crx* and *Otx2* following treatment with the Sstr2 agonist. Treating with reduced concentrations of Met5-enkephalin and naltrexone failed to reproduce the effect seen at saturating concentrations (Fig 2E, F, S2G-J). However, treating with reduced concentrations of the Sstr2 agonist reduced photoreceptor marker gene expression in a dose-dependent manner, suggesting that it was indeed acting through a specific receptor (Fig 2G, S2C-E). Moreover, treatment with excess Sstr2 agonist had no significant effect on *Crx* and *Otx2* expression in explants obtained from *Sstr2^-/-^* mice as measured by t-test, demonstrating that these effects are selectively mediated by activation of Sstr2 signaling (Fig 2H, S2F).

### Sstr2 signaling inhibits photoreceptor specification in embryonic retinal explants

It is unclear whether Sstr2 signaling actually inhibits photoreceptor generation or reduces expression levels of *Otx2* and *Crx*. While qPCR cannot distinguish between these possibilities, scRNA-seq can readily do so. We applied MULTI-seq^40^ to retinal explants grown in a range of Sstr2 agonist concentrations. MULTI-seq allows us to generate a single expression library with cells labeled for each treatment condition (Fig 3A). After clustering and cluster marker identification, we identified the cell type corresponding to each cluster (Fig 3B, S3A). With the sample and cell type identity known for each cell, we quantified and compared the proportions of cell types across the different treatment conditions (Fig 3C). Increasing the concentration of Sstr2 agonists decreased the proportion of photoreceptors and increased the proportion of primary progenitors in the explants. Cell type proportions were significantly different by chi-squared test (p<2.2 E-16). In addition, the level of photoreceptor marker gene expression in photoreceptor clusters was not significantly changed in response to different agonist concentrations by Wilcoxon rank sum test (Fig 3D, S3B). These results demonstrate that, in retinal explants, Sstr2 signaling inhibits photoreceptor specification while increasing the relative fraction of primary RPCs.

**Figure 3.**
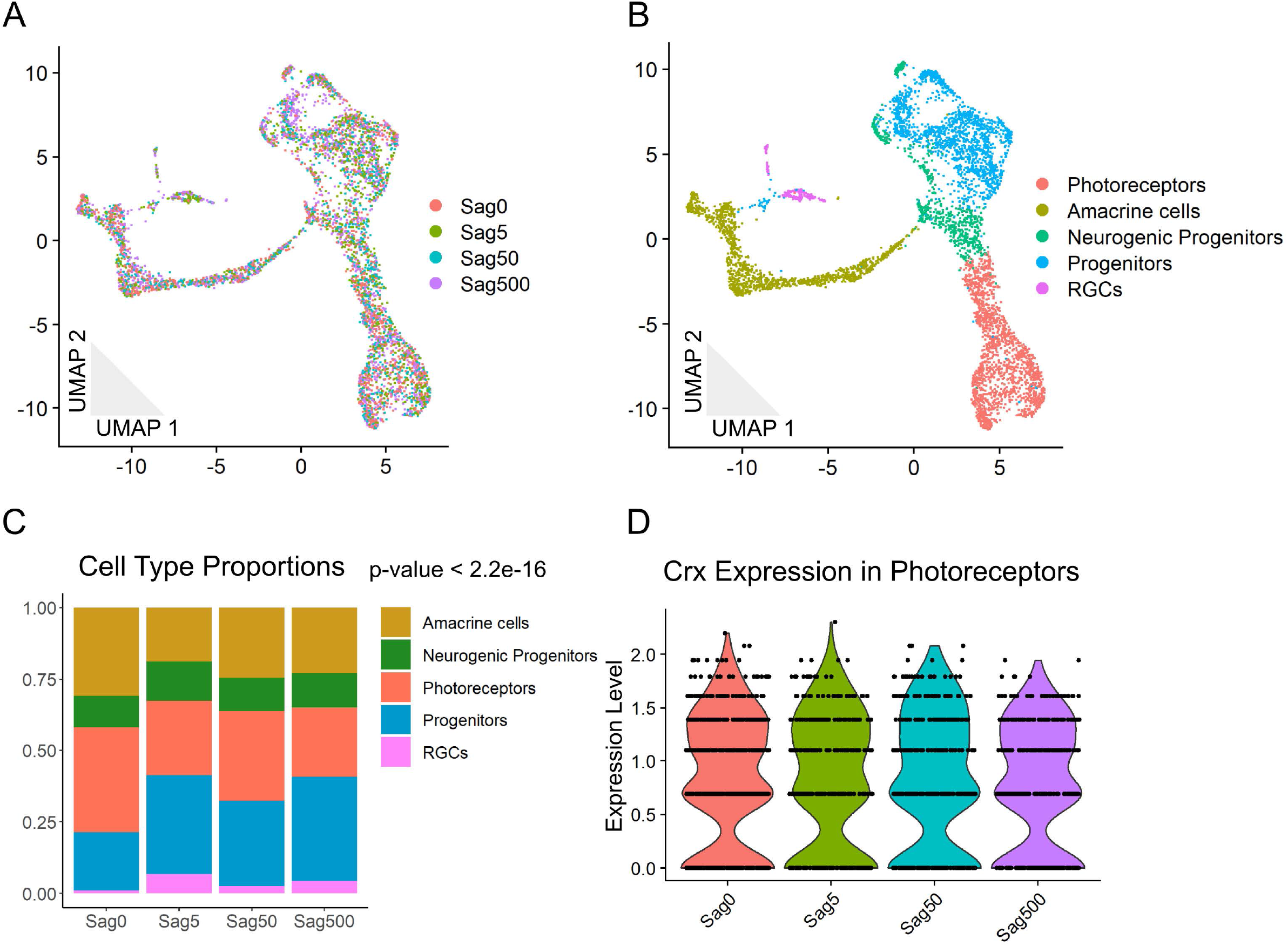
MULTI-seq analysis of explants treated with four different concentrations of Sstr2 agonists reveals changing cell type proportions with increasing treatment. Four concentrations tested: 0, 5 nM, 50 nM, 500 nM Sstr2 agonist. N= 2 samples per condition (except 0, where N=1). N= 5,607 cells. (A) 2D UMAP dimension reduction representation of the pooled explant cells colored by treatment condition. (B) Cells colored by identified cell type. (C) Bar graph representation of the proportion of each cell type within each condition. P-value for significance in difference of proportions calculated using Pearson’s Chi-squared test. (D) Violin plot of expression of *Crx* in photoreceptors for each condition. Was not found to be significantly different by Seurat function ‘FindAllMarkers’.

### No change in cell types is detected in Sstr2^-/-^ retinas

To test whether Sstr2 signaling regulates photoreceptor differentiation *in vivo*, we applied MULTI-seq to the retinas of littermates that were wild type, heterozygous, and homozygous for a targeted deletion of *Sstr2*. We hypothesized that KO mice would have greater proportions of photoreceptors than their littermates. Mice were collected at P0 to match the age at which the explants were tested and P14 to determine whether terminal differentiation and/or survival of specific retinal cell types were affected. Following the same procedure for the sample and cell-type identification as we used for the explants (Fig 4A, B, S4A, B, D, E), we found no significant difference in cell-type proportions between littermates of different genotypes at P0 by chi-squared test (Fig 4C). The genotypes at P14 had significantly different cell type proportions, but this was likely driven by the over enrichment of rods in the Het cells compared to both WT and KO genotypes (Fig S4C). However, *Sstr2* mRNA levels were increased in neurogenic RPCs in *Sstr2* mutants relative to controls as measured by Wilcoxon rank sum test (p-value = 3.35E-5) (Figure 4D), implying that Sstr2 signaling may negatively regulate *Sstr2* transcription.

**Figure 4.**
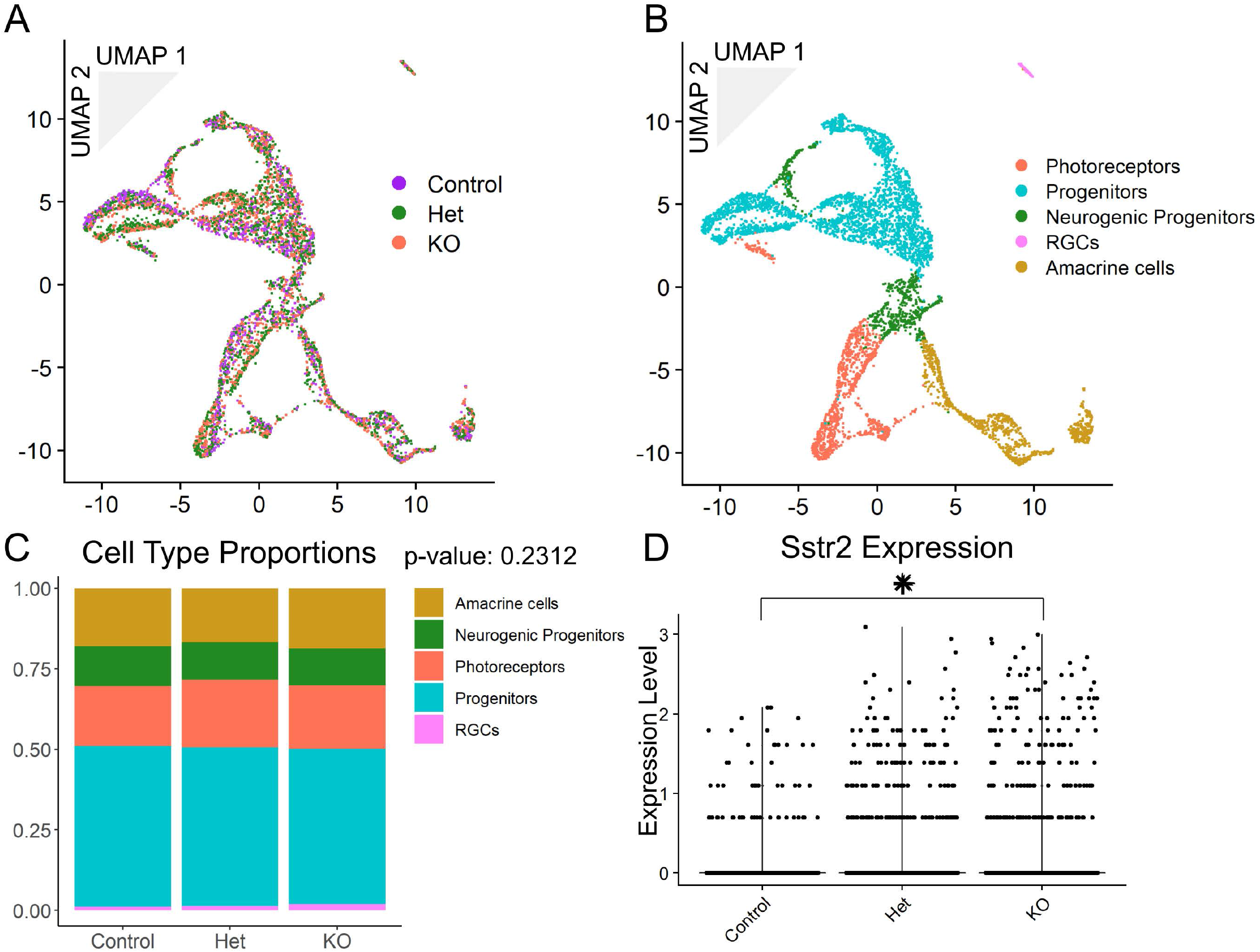
MULTI-seq analysis of wild type, *Sstr2*^+/-^, and *Sstr2-/-* littermates reveals no change in cell-type proportions at P0. N= 2 samples per genotype (except wild type, where N=1). N= 5,706 cells. (A) 2D UMAP dimension reduction representation of the pooled retinal cells colored by genotype. (B) Cells colored by identified cell types. (C) Bar graph representation of the proportion of each cell type within each genotype. P-value for significance in difference of proportions calculated using Pearson’s Chi-squared test. (D) Violin plot of expression of *Sstr2* in cells for each condition. Was found to be significantly different by Seurat function ‘FindMarkers’.

### Sstr2 signaling downregulates genes involved in differentiation and development

We next examined our scRNA-Seq data in more detail to determine which genes showed altered expression in *Sstr2^-/-^* retinas. We hypothesized that though we did not see a change in cell-type proportions *in vivo* that were complementary to those seen in agonist-treated *ex vivo* retinal explants, we might nonetheless observe complementary changes in gene expression. We compared differentially expressed genes from wild type vs. *Sstr2^-/-^* neurogenic RPCs *in vivo* and 0 nM versus 500 nM Sstr2 agonist-treated neurogenic cells *ex vivo*. We identified only one differentially expressed gene in common between these two comparisons. This was *Xist*, and likely simply reflected differences in the sexes of the animals among the different samples. In total, only eleven genes were identified *in vivo* and two identified *ex vivo* by Wilcoxon rank sum test (p< 5E-5) (Suppl. Table 1, 2). Neither of these lists of genes suggested a clear transcriptional impact from Sstr2 perturbation.

We then looked for a mechanism of action for Sstr2 signaling by testing whether genes that differ between genotypes *in vivo* could distinguish cells from different treatments *ex vivo*, as a more nuanced way to evaluate differences in gene expression. To do this, we integrated our *ex vivo* and *in vivo* datasets (Fig 5A, B) and, reasoning that the cells that express *Sstr2* are the most likely to be affected by differences in Sstr2 signaling, selected neurogenic cells that express *Sstr2* from cluster 11 (Fig 5C, D). We then used the 58 genes that are differentially expressed between Sstr2-positive neurogenic cells from wildtype, *Sstr2^+/-^*, and *Sstr2^-/-^* mice to guide the pseudotime program Monocle 2^41^ in aligning all of the Sstr2-positive neurogenic cells along a pseudotime trajectory (Fig 5E, Suppl. Table 3). This analysis identified an unbranched trajectory. While the wild type and heterozygous cells were distributed equally along this trajectory, *Sstr2^-/-^* cells were significantly comparatively enriched at lower pseudotime values (one-sided KS test, p value=0.0213). Interestingly, explant cells treated with lower levels of Sstr2 agonist also appeared to be enriched at lower pseudotime values compared to those treated with the highest concentration of agonist, although this effect did not reach significance (one-sided KS test, p=0.1013). When comparing all low Sstr2 activation conditions (KO retinal and 0, 5, 50 nM agonist explant cells) with all high Sstr2 activation conditions (Het and WT retinal and 500 nM agonist explant cells), cells from low Sstr2 activation conditions were significantly enriched at lower pseudotime values compared to those with higher levels of Sstr2 activation (one-sided KS test, p= 0.0054). None of the samples combined in the low or high groups had significantly different pseudotime distributions as measured by a two-sided KS test. These results led us to conclude that our Sstr2-related perturbations created some shared differences across the two systems.

**Figure 5.**
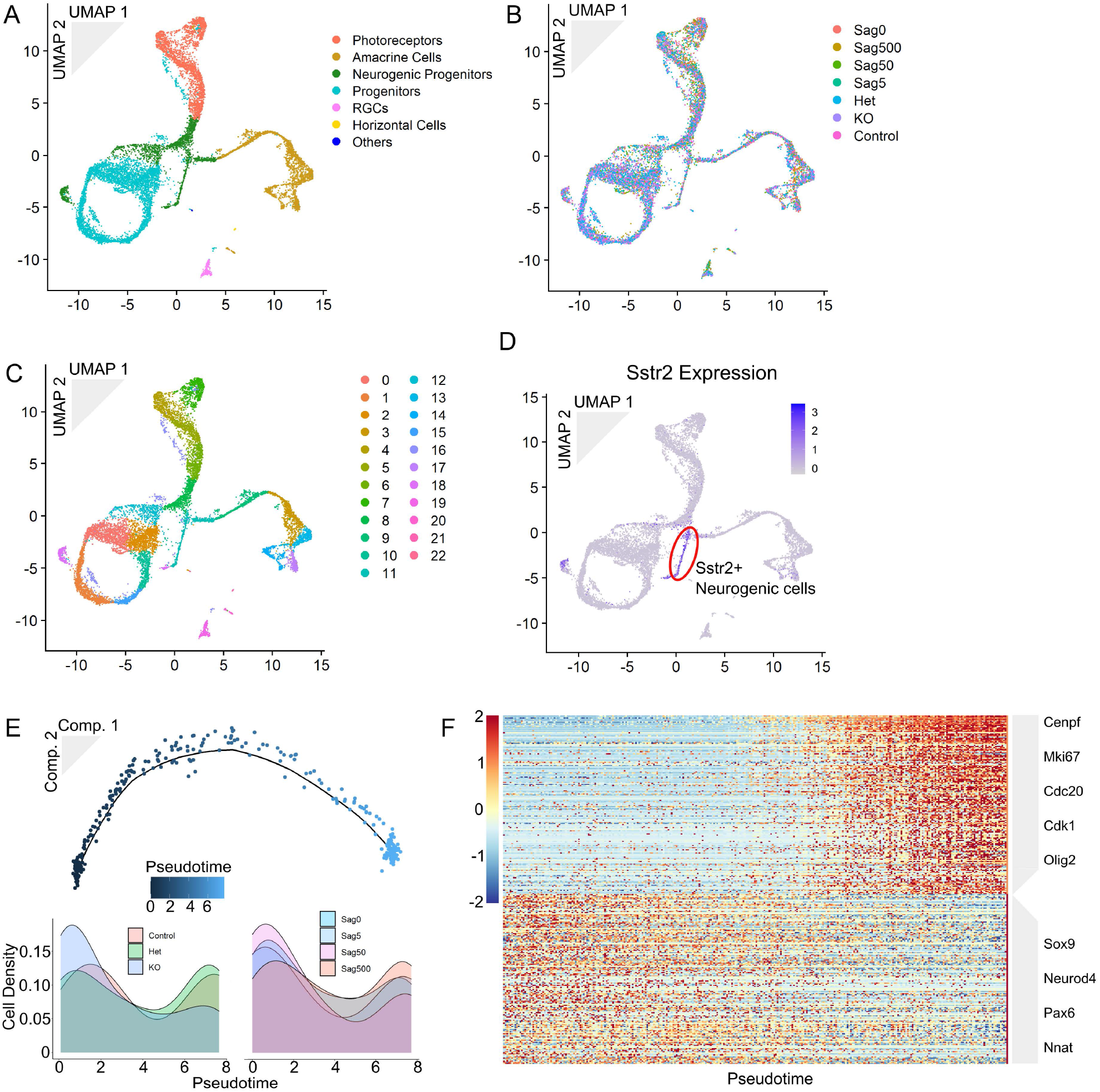
Integrated dataset of explant and retina cells reveals shared genetic changes. The explant and P0 datasets were integrated following the Seurat protocol. N= 11,313 cells. (A) 2D UMAP dimension reduction representation of the integrated datasets colored by identified cell type. (B) Cells colored by genotype or treatment condition. (D) Cells colored by Seurat-identified cluster. (E) Cells colored by Sstr2 expression. (E) Monocle-generated trajectory of *Sstr2+* neurogenic cells from cluster 11 (top). Density plot for cells of each *in vivo* genotype aligned along pseudotime (bottom left). Density plot for cells of each *ex vivo* treatment aligned along pseudotime (bottom left). (F) Heatmap of two clusters of genes that either increase with pseudotime (top 170 genes) or decrease with pseudotime (bottom 161 genes). Genes identified with Monocle 2 function ‘differentialGeneTest()’, q-value cutoff 0.05. Genes clustered by k-means (seed =37).

We next tested for genes that changed in expression across our pseudotime trajectory to see if we could resolve which genes are impacted most by Sstr2 signaling. We identified 331 genes, 170 of which increased with pseudotime and Sstr2 signaling, and 161 of which decreased by likelihood ratio test (q-value<0.05, Fig 5F, Suppl. Table4, 5). The online gene set enrichment tool DAVID^42^ showed a strong enrichment for cell-cycle genes among the genes that increased with pseudotime, including *Cdc20, Cenpf*, and *Mki67*. Genes that regulate neuronal development and differentiation showed decreased expression with pseudotime, including *Neurod4, Nnat*, and *Sox9*. It appears that the pseudotime trajectory captures the transition from proliferative neurogenic RPCs to differentiating post-mitotic neural precursors and that genotypes and conditions with no or low Sstr2 signaling lead to increased cell cycle exit and neurogenesis. In combination with our results obtained from retinal explants, where increasing Sstr2 activation reduced photoreceptor generation, we conclude that Sstr2 signaling inhibits retinal neurogenesis.

## Discussion

We identified four neuropeptides that are transiently, strongly, and specifically expressed during mouse retinal development by analyzing scRNA-seq data obtained from developing mouse retina. We tested whether they regulate retinal development using a combination of small-molecule ligands and qRT-PCR analysis of major retinal cell type markers in embryonic retinal explants. Agonists and antagonists of these neuropeptides—Npy, Gal, and Penk — did not produce significant dose-dependent changes in any retinal cell type markers tested. These results agree with Brodie-Kommit et al., who found no effect on RGC density in Gal-deficient mice^39^. However, agonist-dependent activation of the Sst receptor Sstr2 inhibited photoreceptor generation and led to a compensatory increase in the proportion of primary progenitors. The specificity of these effects was confirmed by their loss in *Sstr2*-deficient retinal explants. However, while Sstr2 signaling inhibits photoreceptor specification in retinal explants, this was not seen *in vivo*, with relative numbers of major retinal cell types unchanged in the *Sstr2* mutant relative to wild type. ScRNA-Seq analysis revealed that retinal explants treated with Sstr2 agonists showed changes in gene expression in neurogenic RPCs that were complementary to those seen in *Sstr2* mutants. These gene expression changes suggest that RGCs may secrete Sst as a quorum signal to negatively regulate retinal neurogenesis. However, the lack of impact on cell type proportion in intact retinas also suggests that Sstr2 signaling is dispensable for regulation of photoreceptor specification *in vivo*.

Similar impacts on retinal cell type proportions have been found with Sonic Hedgehog (Shh) signaling. Wang et al. found that ablation of embryonic RGC-derived Shh signaling in the mouse retina results in precocious cell cycle exit and differentiation of progenitor cells. This resulted in accelerated generation of photoreceptors, depletion of the progenitor pool, and reductions in the proportions of later-born cell types^43^. This matches the phenotype seen in our embryonic retinal explants where treating with an Sstr2-specific agonist resulted in reduced photoreceptor proportion and an increase in the number of primary progenitors. Wang et al. also found that ablation of Shh signaling resulted in a decrease of *Ccnd1* and *Hes1* gene expression, while RPC-enriched genes such as *Hes2* and *Ccnb1* and other cell cycle genes were also decreased upon disruption of Sstr2-dependent signaling. Though we did not observe changes in cell composition in *Sstr2* mutant retinas *in vivo*, pseudotime analysis revealed that neurogenic progenitors in these mutants were relatively more mature than controls, while neurogenic progenitors were relatively less mature in explants treated with Sstr2 agonists. Sstr2+ neurogenic cells in both intact *Sstr2^-/-^* mutant retina and in explants treated with no or low levels of Sstr2 agonist showed increased expression of genes associated with terminal differentiation relative to wildtype, *Sstr2^+/-^*, or explant Sstr2+ neurogenic cells that were treated with high levels of Sstr2 agonist, which in turn showed higher expression of genes enriched in retinal progenitors. Sstr2 signaling may therefore work in tandem with Shh signaling to prevent maturation and differentiation of neurogenic progenitor cells.

A similar dependence on context for the impact of signaling on cell type proportion was reported by Pearson and colleagues^44^. They found that an agonist for the ATP receptor P2RY2 only impacted chick retinal explant development in the absence of the RPE, the endogenous source for ATP in chick retinogenesis. The retinal explants used in the current study lack viable RGCs -- the endogenous source of secreted Sst and Shh at this age. RGCs also express the neuropeptide Pacap, while neurogenic RPCs express the Pacap receptor *Vipr2* during the same interval that *Sst* and *Sstr2* are expressed (Fig S5A, C-E). *Pacap* shows analogous cellular and temporal expression patterns in developing human retina as in mice (Fig S5B-D). We suggest that in the developing retina, RGCs release multiple redundant signals to inhibit cell cycle exit and differentiation in neurogenic progenitors. When retinas are grown as explants, all of the signals are lost, allowing a single exogenous agonist to influence the fate of retinal progenitors. This would explain why Sstr2-mediated signaling dictates cell type proportion *ex vivo* but not *in vivo*.

Interestingly, the proportion of neurogenic progenitors remained consistent across explants at each concentration of Sstr2 agonist, even though fewer cells underwent neurogenesis in explants treated with higher concentrations of agonist. The transition of primary progenitor cells to neurogenic progenitors may therefore be inhibited by the presence of a high concentration of neurogenic cells. If fewer neurogenic cells are exiting mitosis to generate neurons, fewer primary progenitors are entering a neurogenic state. Cell-cell signaling via diffusible factors would be a possible mechanism of action for control of the number of neurogenic progenitors.

We propose that the discovery of an *ex vivo* phenotype that is not maintained *in vivo* and the expression of multiple neuropeptides in RGCs and their receptors in neurogenic progenitor cells describe a complex, redundant system of cell-cell signaling that modulates retinal neurogenesis. Future researchers could interrogate this redundant system to find a role for cell-cell signaling in development *in vivo* through knockout of multiple neuropeptides or receptors. Loss of function of *Vipr2* in combination with *Sstr2* might phenocopy changes in cell-type proportions similar to what is seen with perturbation of Sstr2 signaling *ex vivo* and is seen *in vivo* with loss of *Shh*. We would expect that specifically the cell types produced during the interval in which *Sstr2* and *Vipr2* are expressed would be increased in proportion relative to later-born cell types. This study demonstrates that though *in vivo* experiments are the gold standard for biological relevance, the lack of replication of *ex vivo* results in a more native context does not necessarily equate to a null result.

## Methods

### Retinal Development scRNA-Seq analysis

Sequences and metadata for the mouse (accession number GSE118614) and human (GSE138002) datasets were downloaded from the Gene Expression Omnibus and used to create new Seurat objects following the Seurat SCT workflow^45,46^. Cell type classifications and ages were taken from the downloaded metadata, and used to query the expression of various neuropeptides and receptors. Correlation of neuropeptide expression and each cell type was calculated using the base R function ‘cor()’. Heatmaps were generated with R package pheatmap^47^.

### Mice

Timed pregnant mice were ordered from Charles River Laboratories for the embryonic retinal explant pharmacologic screen. A breeding pair of *Sstr2^+/-^* (C57BL/6NCrl-*Sstr2^em1(IMPC)Mbp^/Mmucd*) mice were ordered from the Mutant Mouse Resource and Research Centers (Davis, CA) and outbred to CD1 mice. The *Sstr2^+/-^* F1 progeny were bred to produce the *Sstr2^+/+^, Sstr2^+/-^*, and *Sstr2^-/-^* mice used in this study. Similarly, two Sst^+/-^ male mice^48^ were received from Malcolm Low at the University of Michigan and outbred to CD1 mice. The Sst^+/-^ F1 progeny were then bred to produce the *Sst^+/+^, Sst^+/-^*, and *Sst^-/-^* mice used in this study. The animals were cared for and euthanized in compliance with Johns Hopkins IACUC guidelines.

### Retinal explant culture

Retinas were dissected from embryonic day (E)14 or E18 embryos from timed pregnant mice, age verified by crown-rump length. Dissected retinas were then flattened using a series of radial cuts, and mounted on 0.2 uM Nuclepore Track-Etch membranes on 2 ml DMEM F12, 10% FBS with 0.1% streptomycin/puromycin in 12-well plates. The explants were incubated at 37 degrees Celsius, 5% CO2 for 6 days for E14 explants, and 4 days for E18 explants. On even-numbered days, 90% of the explant culture media was replaced with new media. Treatment compounds were included in the explant culture media at the concentrations listed in the text and ordered from Tocris Bioscience.

### Explant RNA extraction and cDNA preparation

Embryonic retinal explants were removed from the nuclepore membrane and media and quick-frozen at −80 C or on dry ice. Total RNA was extracted from explants using the Qiagen miRNeasy Mini Kit with optional DNase digest and drying steps. The concentration of the resultant RNA was determined using a ThermoScientific NanoDrop 1000. 400 ng RNA was reversed transcribed to cDNA using the Invitrogen SuperScript IV First-Strand Synthesis System. The cDNA was diluted to 1 ng/ul in nuclease-free water.

### Explant qPCR analysis

We used the bimake.com 2x SYBR Green qPCR Master Mix. Four ng cDNA was used for each 20 ul reaction. Three technical replicates were assayed for each sample. Samples were run on a standard 2-step amplification program on Applied Biosystems MicroAmp Fast Optical 96-Well or MicroAmp Optical 384-Well reaction plates on an Applied Biosystems StepOne Plus or ViiA 7, respectively. Primer sequences used either had been previously validated in the Blackshaw lab or were taken from the Harvard Primer Bank or generated using Primer3^40,49^. Those generated on Primer3 were required to span an exon-exon junction. All primers were verified to produce a single reaction product with traditional PCR before use for qPCR. All plots generated with ggplot2 R package^50^.

### MULTI-seq

Samples were dissociated using the Worthington papain cell dissociation system. MULTI-seq was performed to pool samples prior to 10x Chromium Single Cell 3’ v3.0 GEM generation and library prep following the Gartner lab’s protocol using lipid-tagged oligos provided by the Gartner lab. Quality control and sequencing were performed by the Johns Hopkins Transcriptomics and Deep Sequencing Core.

### scRNAseq analysis

Sequencing results were processed through the 10x CellRanger pipeline and the output loaded into Seurat^45^. Sample of origin for each cell was determined using the deMULTIplex R package out of the Gartner lab^40^. Seurat objects were analyzed following the Seurat SCT workflow. Cell type was determined cluster by cluster based on marker gene expression derived from the Seurat function ‘FindAllMarkers’ using default parameters^45^. This function was also used to identify differential gene expression across conditions/genotypes. Alternatively, and where noted in Results, the Seurat function ‘FindMarkers’ was used to find differential expression between two groups of cells.

Seurat objects were integrated following the Seurat Integration and Label Transfer workflow. The pseudotime trajectory was constructed and differential genes identified using the semi-supervised Monocle 2 workflow^41^. Density plots generated using ggplot2. Differential genes were clustered using the base R function ‘k-means’, and heatmap was generated using pheatmap.

## Acknowledgments

This work was supported by a grant from the NIH (R01EY020560) to S.B, the Maryland Stem Cell Research Fund (2019-MSCRFF-5124) to DWK, and an award from the Visual Neuroscience Training Program to K.W. We thank Transcriptomics and Deep Sequencing Core (Johns Hopkins) for sequencing the scRNA-Seq libraries. Sst^+/-^ mice were provided by Malcolm Low at the University of Michigan.

## Contribution

S.B. conceived and initiated the study. K.W. performed all qRT-PCR, scRNA-Seq and histological analysis. D.W.K. helped analyze scRNA-Seq data. All authors wrote the paper.

## Data availability

All sequencing data will be available on GEO upon publication.

## Supplemental figures

**Figure S1.**
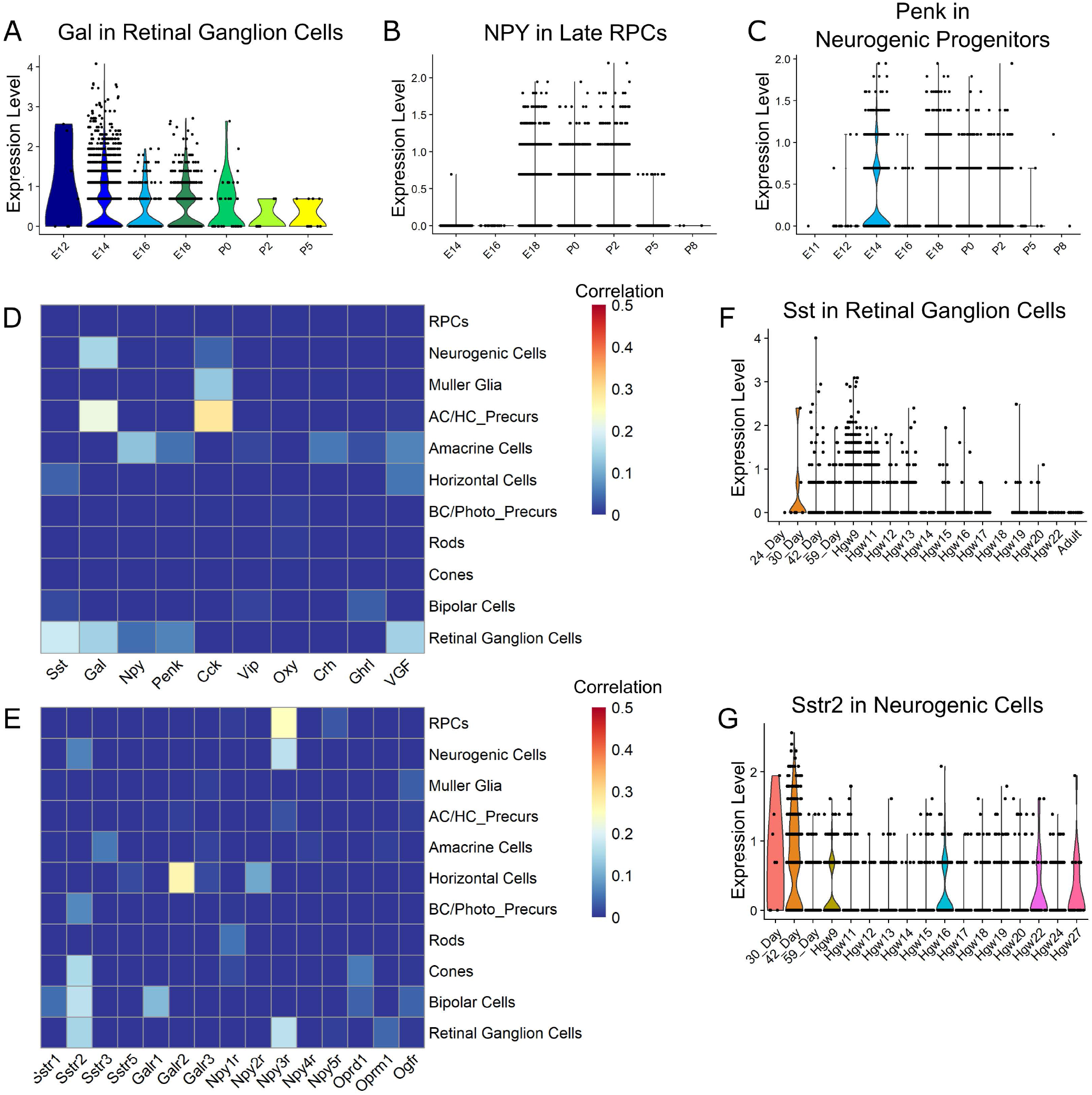
Cell-specific expression of neuropeptides and neuropeptide receptors in developing human retina. (A) Violin plot of expression of galanin in retinal ganglion cells across development. (B) Violin plot of expression of neuropeptide Y in late progenitor cells across development. (C) Violin plot of proenkephalin expression in neurogenic progenitor cells across development. (D) Heatmap of correlation of expression of a neuropeptide with cell type in human retinal development. (E) Correlation of expression of a receptor with cell type in human retinal development. Sstr4 was not expressed at detectable levels in the dataset. (F) Violin plot of expression of Sst in retinal ganglion cells across human development. (G) Violin plot of expression of Sstr2 in neurogenic progenitor cells in human retinal development. Correlation calculated using base R function ‘cor()’.

**Figure S2.**
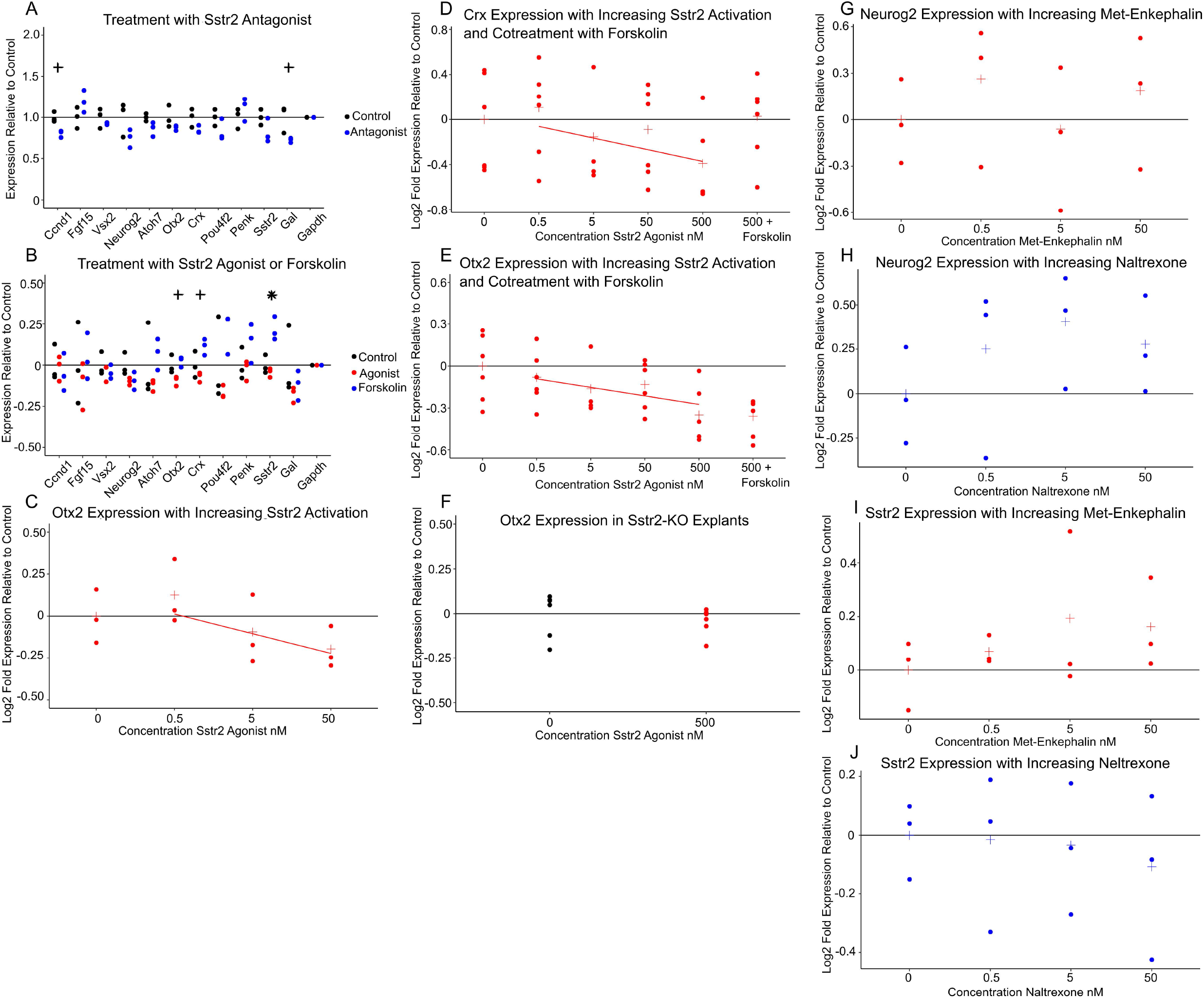
Treatment of embryonic retinal explants with neuropeptides, agonists and antagonists produces less impressive effects on cell type marker genes and signals. (A) Expression relative to control of marker genes for explants treated with Sstr2 antagonist CYN 154806. Concentration 1 uM. N=3 each condition. Explants treated E14-P0. (B) Marker gene expression relative to average control of explants treated with forskolin or Sstr2 agonist. Concentration 500 nM. N=3 each condition. Explants treated from E14-P0. (C) *Otx2* expression in explants treated with increasing levels of Sstr2 agonist. N=3 each condition. Explants treated E14-P0. (D) *Crx* expression in explants treated with increasing levels of SSTR2 agonist or 500 nM agonist and 500 nM Forskolin. N=6 each condition (except 5 nM treatment where N=4). Fitted line added for emphasis. (E) Otx2 expression in explants treated with increasing levels of Sstr2 agonist or 500 nM agonist and 500 nM Forskolin. N=6 each condition (except 5 nM treatment where N=4). Fitted line added for emphasis. (F) *Otx2* expression relative to average control of Sstr2 -/- explants treated with 500 nM Sstr2 agonist. N=6 each condition. (G) Neurog2 expression relative to average control in explants treated with increasing concentration of Met-Enkephalin. N=3 each condition. (H) *Neurog2* expression relative to average control in explants treated with increasing concentration of naltrexone. N=3 each condition. (I) Sstr2 expression relative to controls in explants treated with increasing concentration of Met-enkephalin. N=3 each condition. (J) *Sstr2* expression relative to average control in explants treated with increasing concentration of Naltrexone. N=3 each condition. P-value calculated using ANOVA. + indicates a nominally significant p-value below 0.05. * indicates a p-value significant at a Bonferroni correction below 0.0045

**Figure S3.**
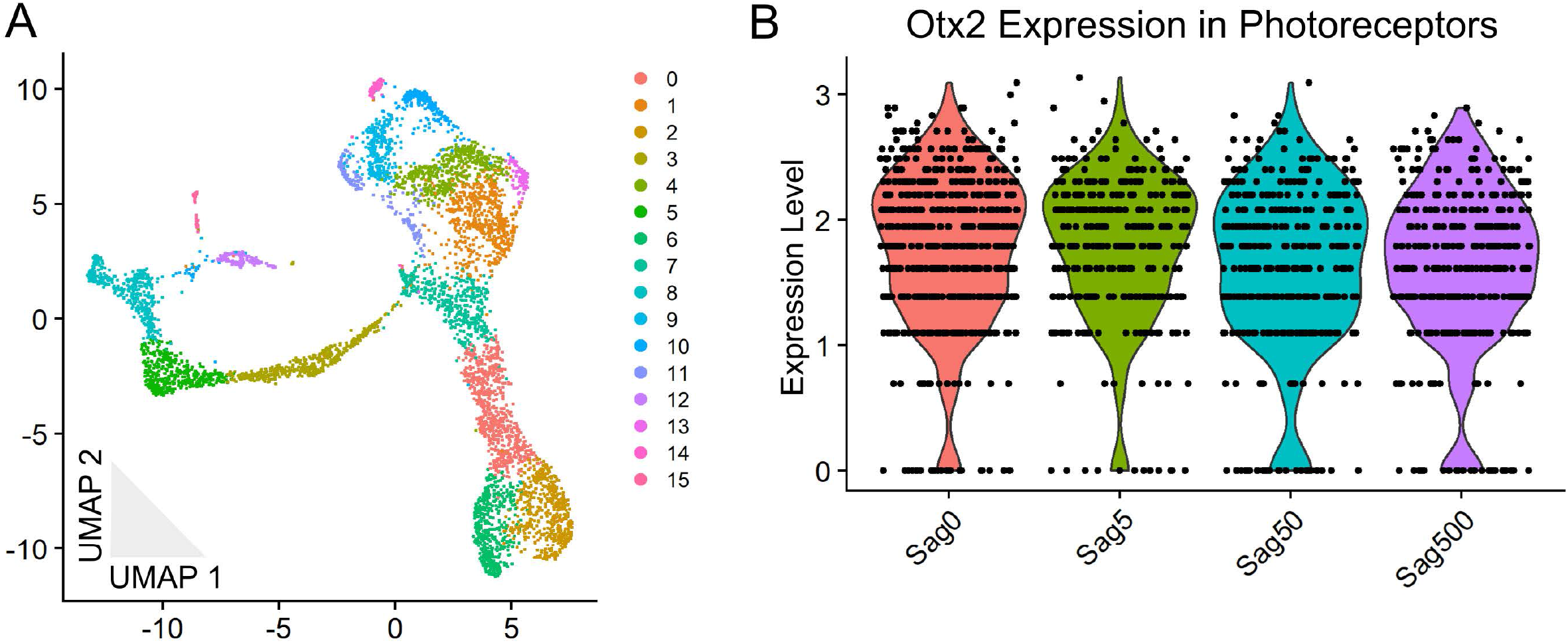
MULTI-seq analysis MULTI-seq analysis of explants treated with four different concentrations of Sstr2 agonist reveals changing cell type proportions with increasing treatment. Four concentrations tested: 0, 5 nM, 50 nM, 500 nM Sstr2 agonist. N= 2 samples per condition (except 0, where N=1). N= 5,607 cells. (A) 2D UMAP dimension reduction representation of the pooled explant samples colored by Seurat-identified cluster. (B) Violin plot of expression of *Otx2* in photoreceptors for each condition. Was not found to be significantly different by Seurat function ‘FindAllMarkers’.

**Figure S4.**
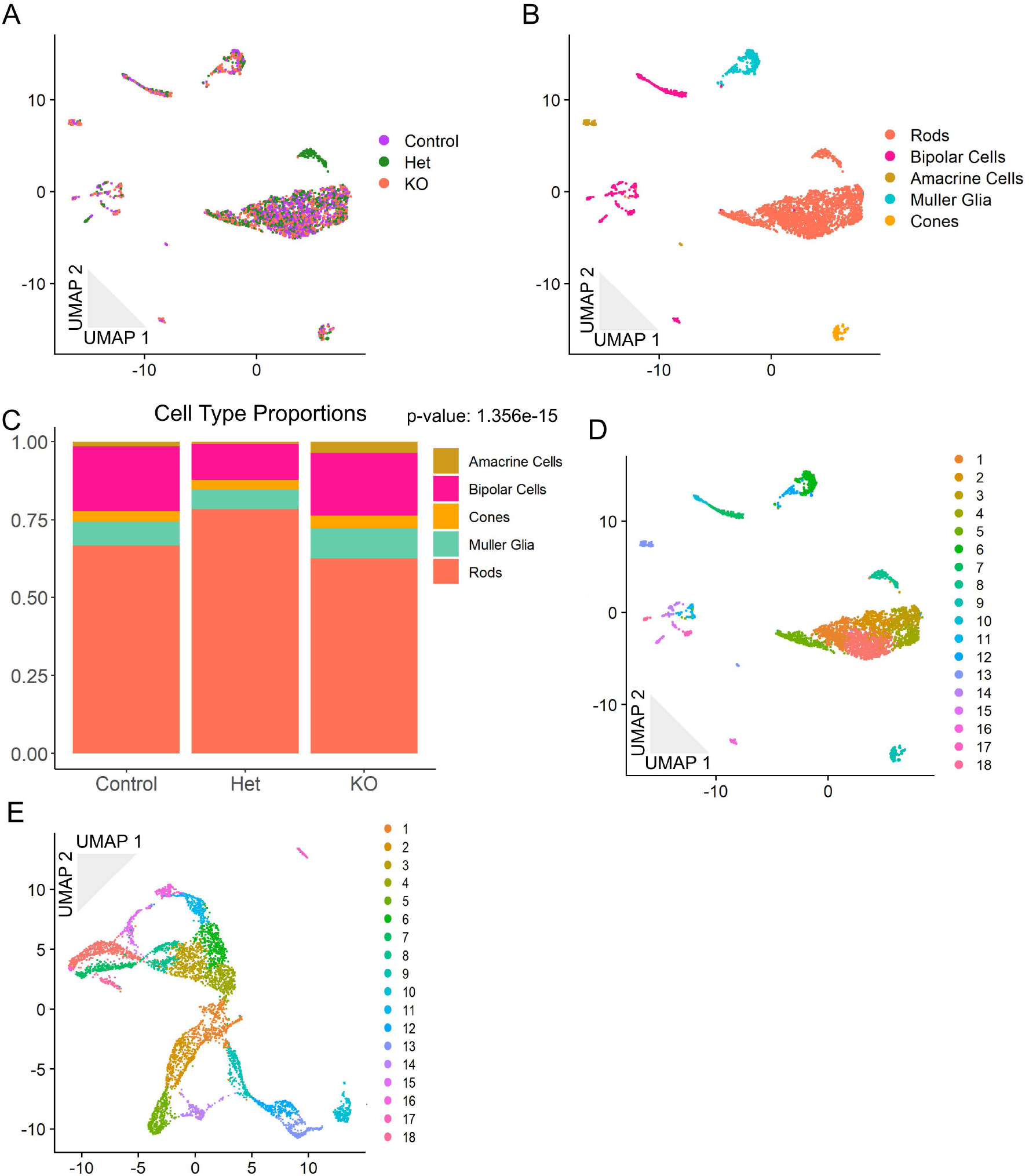
MULTI-seq analysis of wild type, *Sstr2*^+/-^, and *Sstr2^-/-^* littermates reveals no change in cell-type proportions at P14. N= 2 samples per genotype (except wildtype where N=1). N= 3,298 cells. (A) 2D UMAP dimension reduction representation of the pooled retinal cells colored by genotype. (B) Cells colored by identified cell types. (C) Bar graph representation of the proportion of each cell type within each genotype. P-value for significance in difference of proportions calculated using Pearson’s Chi-squared test. (D) Cells colored by Seurat-identified cluster. (E) 2D UMAP dimension reduction representation of the pooled P0 samples colored by Seurat-identified cluster.

**Figure S5.**
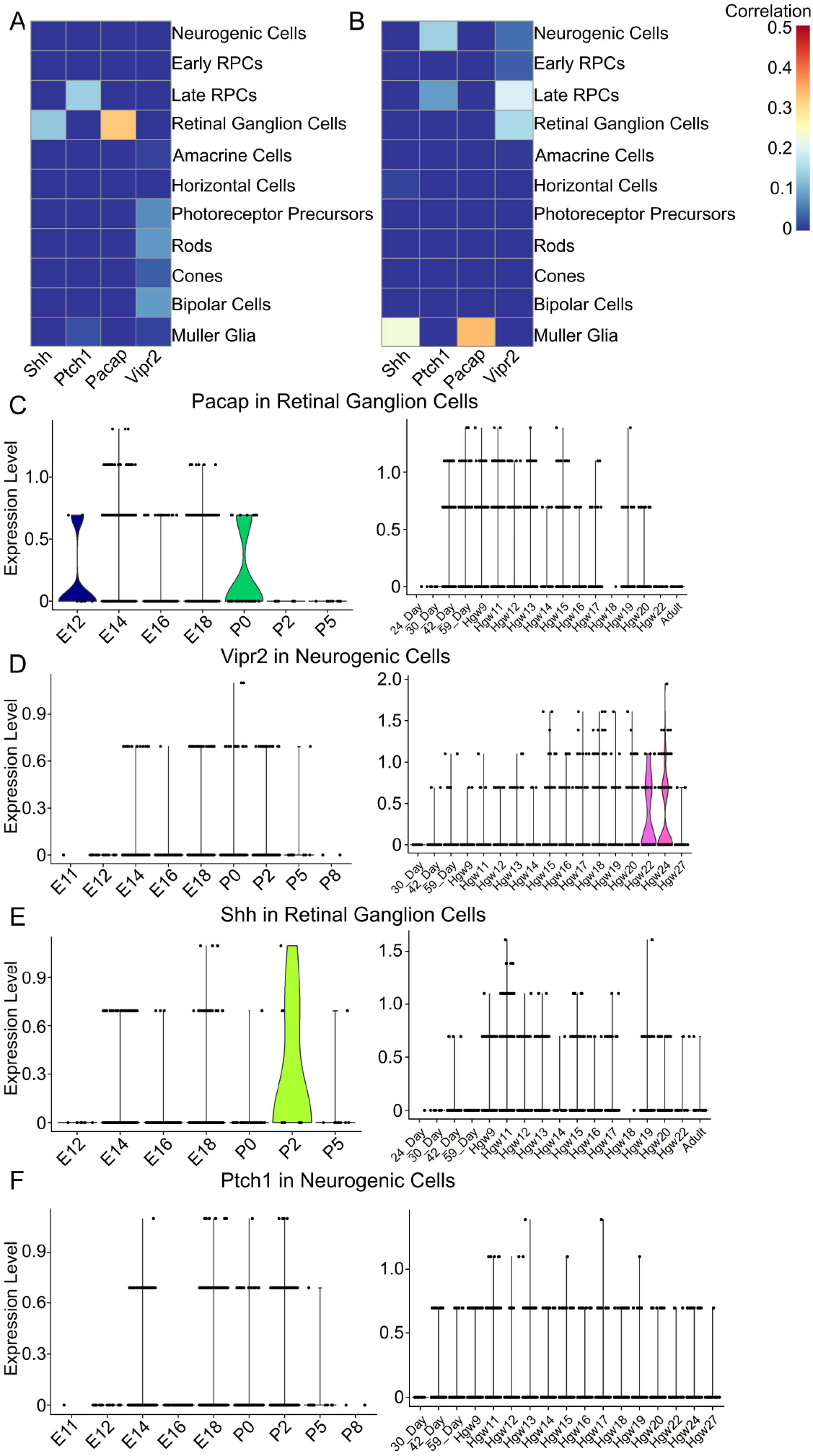
Additional neuropeptides show cell type and time point specificity in mouse and human retinal development. May have a redundant function with Sst. (A) Heatmap of correlation of expression of neuropeptide or receptor with cell type in mouse retinal development. (B) Heatmap of correlation of expression of neuropeptide or receptor with cell type in human retinal development. (C) Violin plots of expression of *Pacap* in retinal ganglion cells across mouse (left) and human (right) retinal development. (D) Violin plots of expression of Pacap receptor Vipr2 in neurogenic progenitor cells across mouse (left) and human (right) retinal development. (E) Violin plots of expression of ATP receptor *P2ry2* in neurogenic progenitor cells across mouse (left) and human (right) retinal development.

## Supplemental tables

**Supplemental Table 1**: Genes differentially expressed in neurogenic RPCs in 0nm vs. 500 nm Sstr2 agonist.

**Supplemental Table 2**: Genes differentially expressed in neurogenic RPCs in wildtype vs. *Sstr2^-/-^* mice.

**Supplemental Table 3:** Genes used for pseudotime analysis of neurogenic RPCs in wildtype, *Sstr2^+/-^* and *Sstr2^-/-^* mice.

**Supplemental Table 4:** Genes increased with pseudotime and higher levels of Sstr2 signaling.

**Supplemental Table 5:** Genes decreased with pseudotime and higher levels of Sstr2 signaling.

